## supplementary material for "Emergence of power-law distributions in protein-protein interaction networks through study bias"

### Supplementary Table legends

Supplementary Table 1: Enrichment results for all the tested annotation categories and for all hub definitions.

### Supplementary Figures

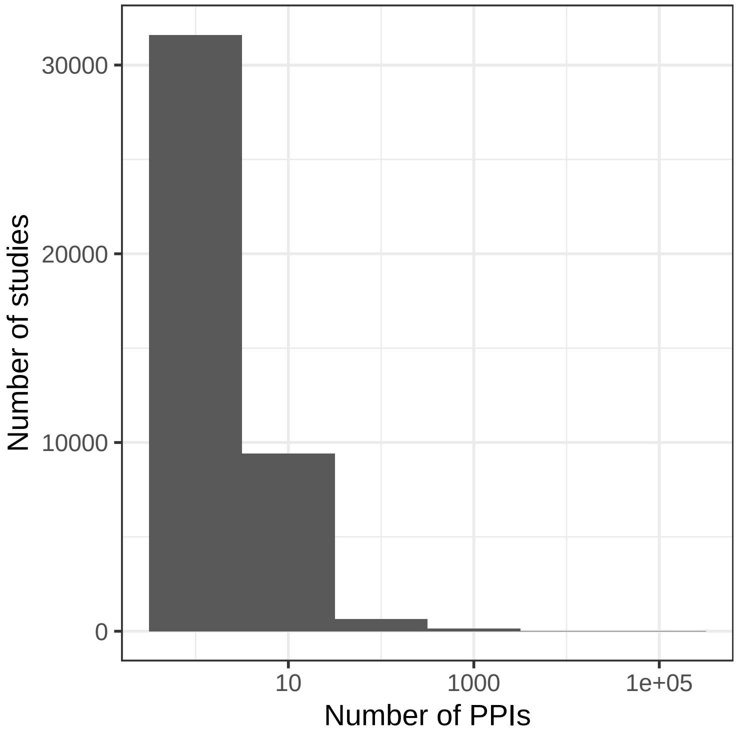


**Supplementary Figure 1.** Histogram of PPI number per study in the aggregated network.

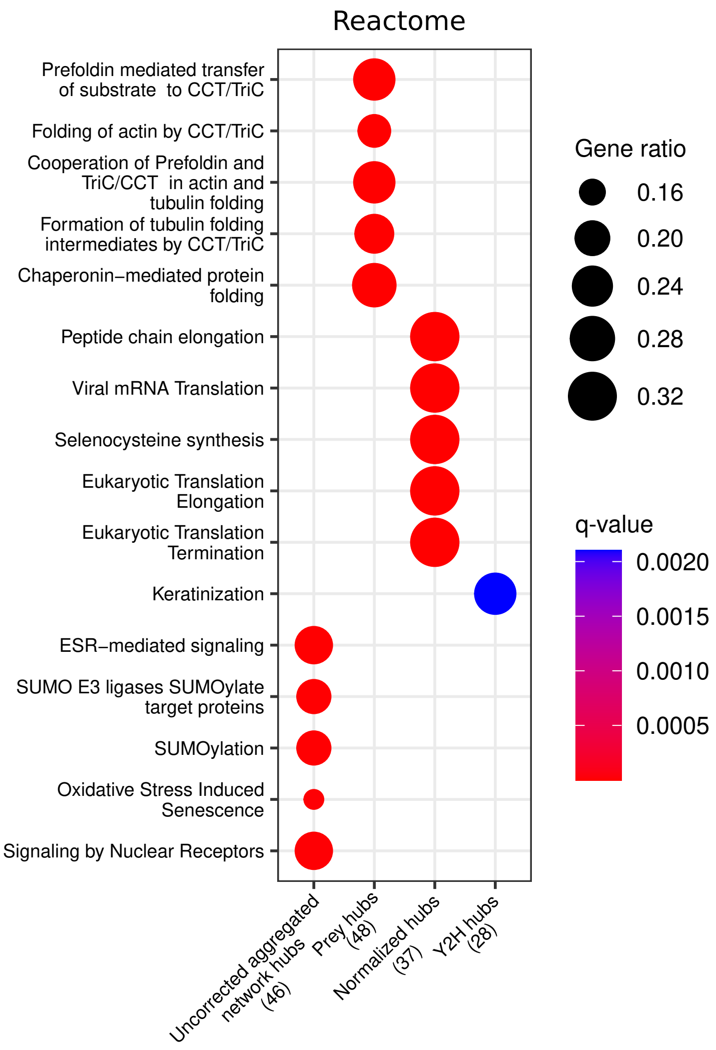


**Supplementary Figure 2.** Reactome enrichment analysis of the top 50 corrected and uncorrected hubs. The numbers in brackets represent the number of hubs included in the reference databases, and the ’Gene ratio’ represents the fraction of hubs included in the corresponding pathway.

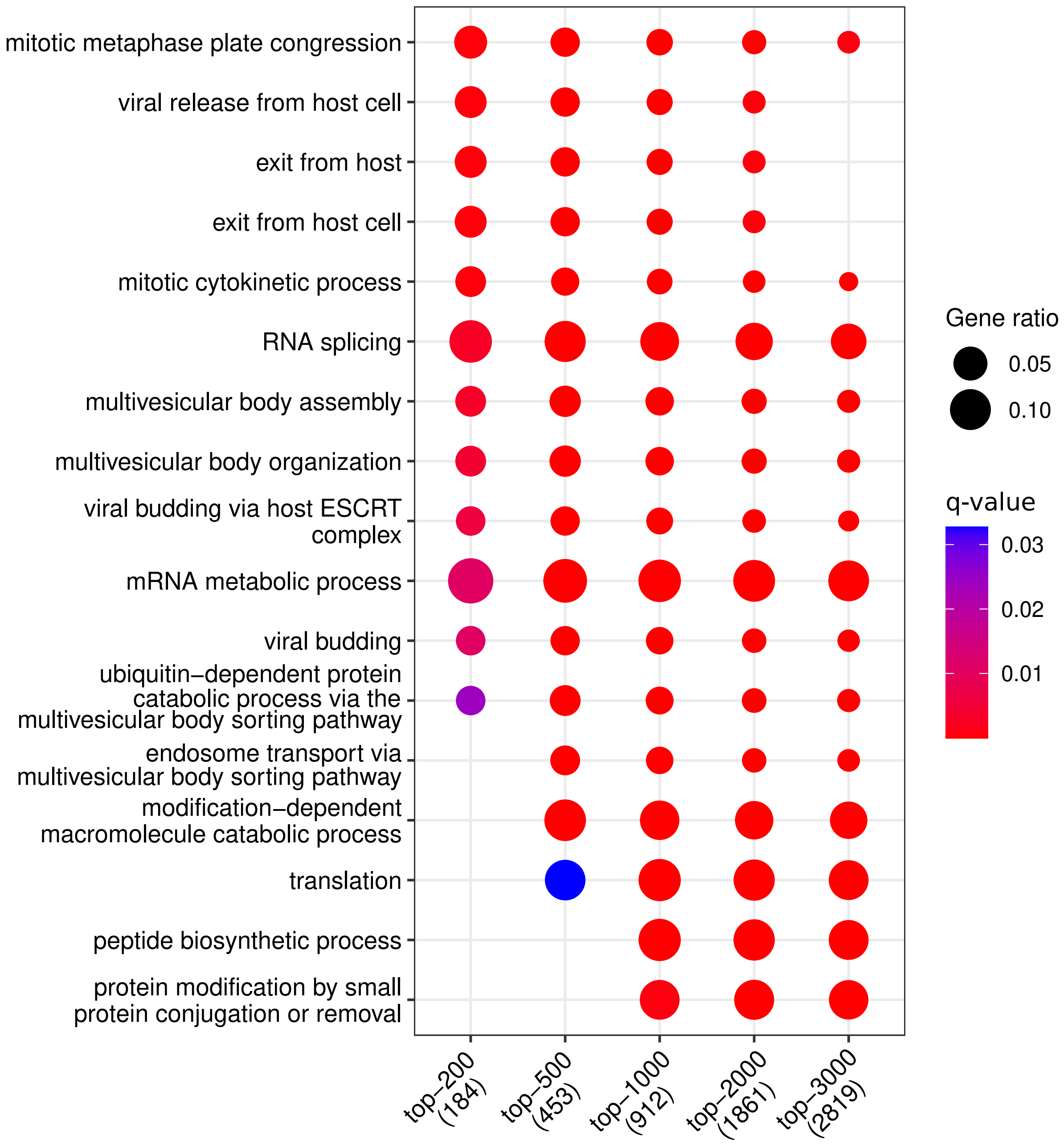


**Supplementary Figure 3.** Gene Ontology enrichment analysis results of the top-200, top-500, top-1000, top-2000, and top-3000 most abundant proteins.

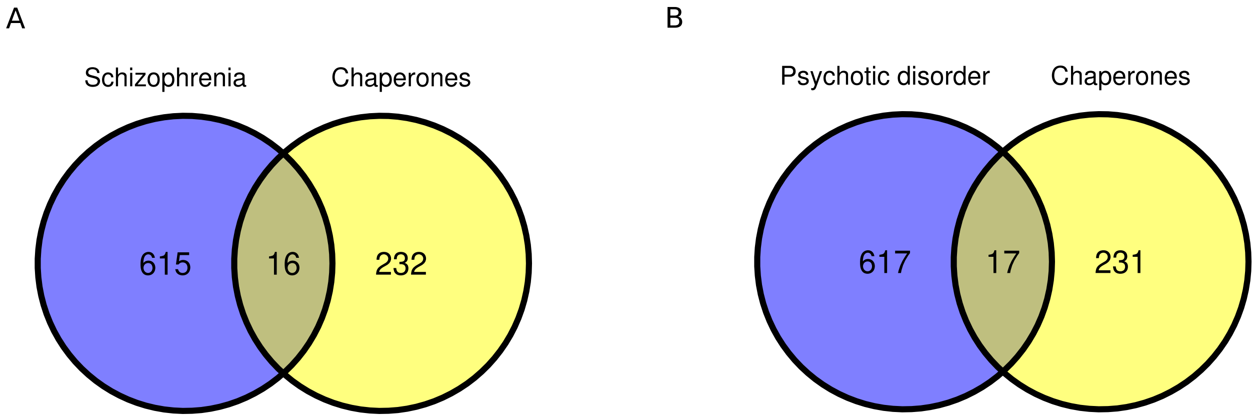


**Supplementary Figure 4.** (A) Overlap between schizophrenia-related genes and chaperones ($P = 0.01$; one-sided Fisher test). (B) Overlap between psychotic disorder-related genes and chaperones ($P = 0.005$; one-sided Fisher test).

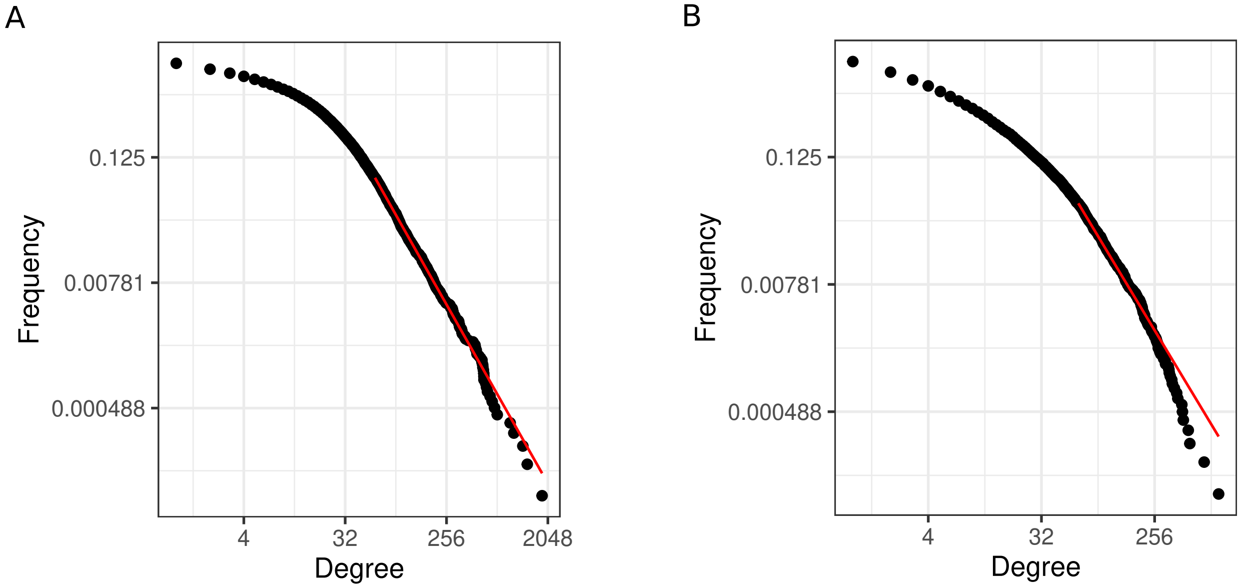


**Supplementary Figure 5.** (A) Degree distribution for observed PPI network obtained via aggregation of all AP-MS studies annotated in IntAct. Node degrees are PL-distributed ($P = 0.34$). (B) Degree distribution for observed PPI network obtained via aggregation of all Y2H studies annotated in IntAct. Node degrees are not PL-distributed ($P = 0.01$).

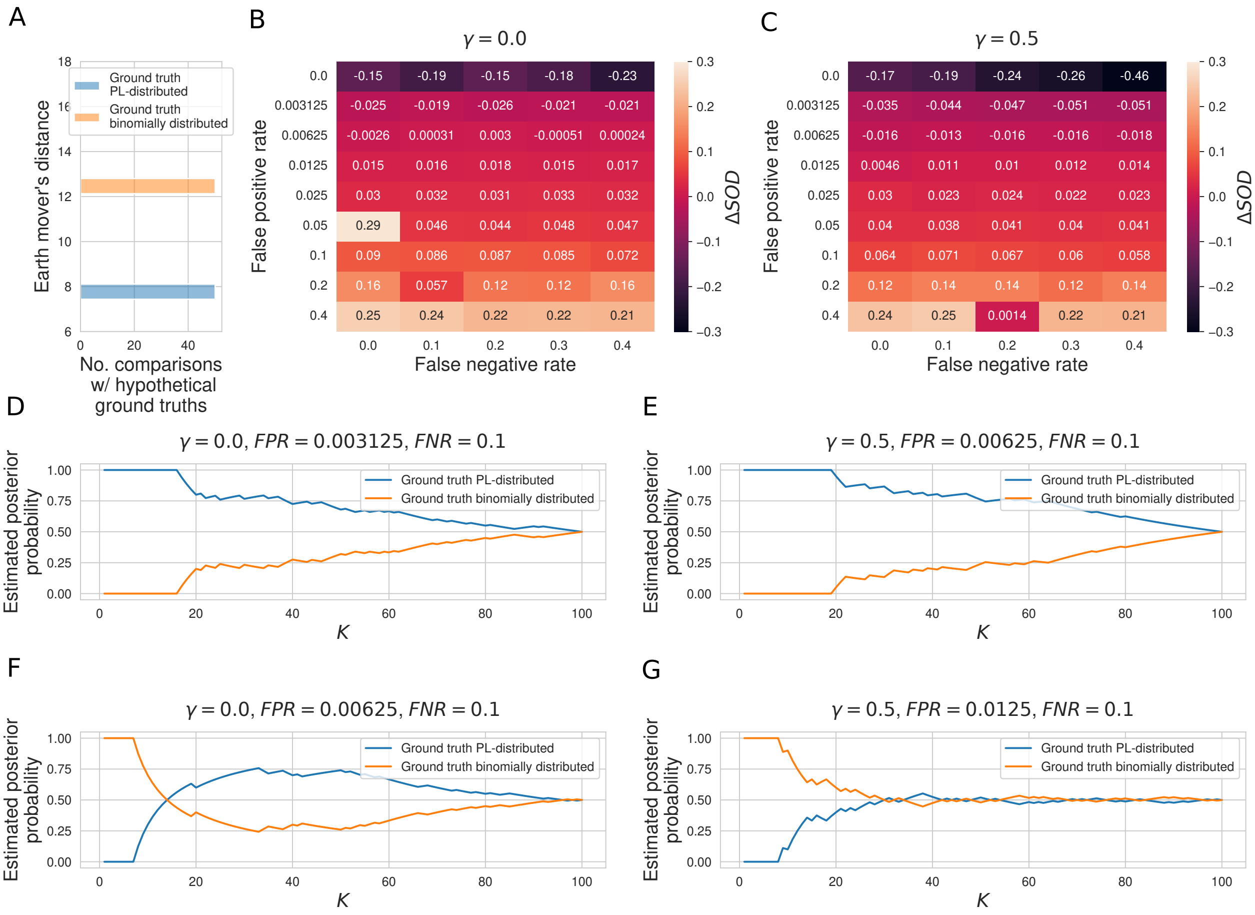


**Supplementary Figure 6.** (A) Histogram of earth mover's distances between the degree distribution of the observed PPI network $G_{\text{IntAct}}$ obtained via aggregation of all Y2H studies annotated in IntAct and the degree distributions of 50 PL-distributed and 50 binomially distributed hypothetical ground truth networks. (B, C) Signed relative differences $\Delta SOD \in [-1, 1]$ between sum of distances between degree distribution of $G_{\text{IntAct}}$ and degree distributions of networks simulated from, respectively, PL-distributed and binomially distributed hypothetical ground truth networks, given different choices of the hyper-parameters $FPR$, $FNR$, and $\gamma$. Negative values of $\Delta SOD$ indicate that $G_{\text{IntAct}}$ is more similar to simulated networks emerging from PL-distributed hypothetical ground truths; positive values are indicative of the opposite scenario. (D–G) Posterior probabilities that $G_{\text{IntAct}}$ emerged from a PL-distributed or from a binomially distributed ground truth interactome, estimated via $K$-nearest neighbors classification. (D, E) Estimated posterior probabilities just before the tipping points in the false positive rate. (F, G) Estimated posterior probabilities just after the tipping points.
